# NREM sleep oscillations and their relations with sleep-dependent memory consolidation in early course psychosis and first-degree relatives

**DOI:** 10.1101/2023.10.30.564703

**Authors:** Dan Denis, Bengi Baran, Dimitrios Mylonas, Courtney Spitzer, Nicolas Raymond, Christine Talbot, Erin Kohnke, Robert Stickgold, Matcheri Keshavan, Dara S. Manoach

## Abstract

Sleep spindles are believed to mediate sleep-dependent memory consolidation, particularly when coupled to neocortical slow oscillations. Schizophrenia is characterized by a deficit in sleep spindles that correlates with reduced overnight memory consolidation. Here, we examined sleep spindle activity, slow oscillation-spindle coupling, and both motor procedural and verbal declarative memory consolidation in early course, minimally medicated psychosis patients and non-psychotic first-degree relatives. Using a four-night experimental procedure, we observed significant deficits in spindle density and amplitude in patients relative to controls that were driven by individuals with schizophrenia. Schizophrenia patients also showed reduced sleep-dependent consolidation of motor procedural memory, which correlated with spindle density. Contrary to expectations, there were no group differences in the consolidation of declarative memory on a word pairs task. Nor did the relatives of patients differ in spindle activity or memory consolidation compared with controls, however increased consistency in the timing of SO-spindle coupling were seen in both patient and relatives. Our results extend prior work by demonstrating correlated deficits in sleep spindles and sleep-dependent motor procedural memory consolidation in early course, minimally medicated patients with schizophrenia, but not in first-degree relatives. This is consistent with other work in suggesting that impaired sleep-dependent memory consolidation has some specificity for schizophrenia and is a core feature rather than reflecting the effects of medication or chronicity.

**Statement of significance:** We investigated sleep neurophysiology and memory consolidation in minimally-medicated, early course psychosis patients and unaffected first-degree relatives of patients in a four-night study. Early-course schizophrenia patients had a sleep spindle deficit that correlated with reduced procedural memory consolidation. Although first-degree relatives did not show any deficit in spindles or memory consolidation, the spindles of both relatives and patients showed increased consistency of the temporal coupling of spindles with slow oscillations compared with controls. These results suggest that sleep spindle and memory consolidation deficits are a core feature of schizophrenia. Spindle deficits may be a biomarker of schizophrenia modifiable by treatment, and motivate the development of targeted novel interventions.

## Introduction

Schizophrenia is characterized by a deficit in sleep spindles that, in chronic patients, generally presents in the context of unaltered sleep quality and architecture (Ferrarelli et al., 2007, 2010; Lai et al., 2021; Mylonas et al., 2020; Wamsley et al., 2012). Sleep spindles, a defining feature of stage 2 non-rapid eye movement sleep (N2), are brief (∼ 1 second) bursts of ∼12-15 Hz synchronous activity initiated by the thalamic reticular nucleus (Fernandez & Luthi, 2020). Sleep spindles are propagated to the cortex, where they induce long-term potentiation (LTP) and synaptic plasticity, facilitating overnight memory consolidation (Peyrache & Seibt, 2020; Rosanova & Ulrich, 2005; Seibt et al., 2017; Steriade & Timofeev, 2003; Werk et al., 2005). Chronic patients with schizophrenia have reduced sleep-dependent procedural and declarative memory consolidation (Baran et al., 2018; Göder et al., 2015; Manoach et al., 2004; see Demirlek & Bora, 2023 for meta-analysis) that correlates with spindle and slow oscillation (SO)-spindle coupling deficits (Göder et al., 2015; Manoach et al., 2010; Wamsley et al., 2012). In young, early course antipsychotic-naïve schizophrenia patients and non-psychotic first-degree relatives, sleep-dependent memory consolidation has not been examined, but spindle deficits correlate with more general cognitive impairments (Manoach et al., 2014). These findings indicate that spindle deficits are unlikely to reflect the effects of medication or chronicity. Instead, they may be a core feature of schizophrenia that is present early in its course and impair cognition. The goals of the present study were to replicate and extend these findings by investigating sleep spindles, slow oscillation (SO)-spindle coupling, and both motor procedural and verbal declarative memory consolidation in early course, minimally medicated psychosis patients and non-psychotic first-degree relatives.

Sleep-dependent memory consolidation relies on the temporal coordination of hippocampal sharp wave ripples (∼80-150 Hz) with sleep spindles and neocortical SOs (∼.5-1.25 Hz) (Klinzing et al., 2019). This inter-regional dialogue is thought to facilitate the transfer of recently encoded memories from temporary indexing in the hippocampus to longer-term distributed storage in the cortex (Helfrich et al., 2019; Klinzing et al., 2019; Latchoumane et al., 2017; Staresina et al., 2015). Evidence that it is SO-spindle coupling that is crucial to memory consolidation comes from studies in healthy participants demonstrating that SO-coupled spindles are the best predictor of consolidation, independent of uncoupled spindles (Denis et al., 2021; Schreiner et al., 2021; Solano et al., 2022). In schizophrenia, SO-spindle coupling has been less frequently assessed. In chronic, medicated patients initial findings reveal that SO-spindle timing is largely spared (Demanuele et al., 2016; Mayeli et al., 2022), though an *increased* consistency in the phase of the SO in which the spindle peak has been observed (Kozhemiako et al., 2022; Mylonas et al., 2020). SO-spindle coupling is also associated with memory consolidation in patients (Demanuele et al., 2016; Mylonas et al., 2020). A recent study on patients at clinical high risk for psychosis revealed no differences in SO-spindle timing compared to matched controls (Mayeli et al., 2022). To our knowledge, there are no previous studies that examined SO-spindle coupling or its relations with memory consolidation in early course patients and first-degree relatives.

Importantly, among psychotic disorders, spindle deficits may be specific to schizophrenia and do not appear to be fully accounted for by medication side effects or disease chronicity (Manoach et al., 2014). Emerging research has documented spindle deficits in early course, antipsychotic naive patients, as well as first-degree relatives of patients (D’Agostino et al., 2018; Kaskie et al., 2019; Schilling et al., 2017), but not in individuals with other psychotic disorders in either early course (Manoach et al., 2014) or chronic (Ferrarelli & Tononi, 2017) samples. These findings suggest that the sleep spindle deficit in schizophrenia is an endophenotype reflecting genetic risk for schizophrenia (Manoach et al., 2016; Schilling et al., 2022). It is not known whether early course patients and unaffected relatives show associated deficits in sleep-dependent memory consolidation.

In the present study, we utilised a multi-night experimental procedure and high-density electroencephalography (EEG) to investigate these gaps in the literature. Specifically, we hypothesised that 1) Early course patients and relatives would show a deficit in sleep spindle density compared to healthy controls. 2) Patients and relatives would also show reduced overnight procedural and declarative memory consolidation. 3) Sleep spindles, particularly those coupled to slow oscillations, would correlate with overnight memory consolidation. We also hypothesised that any deficits in early course patients would be driven by schizophrenia patients compared to individuals with non-schizophrenia psychoses.

## Methods

### Participants

A total of 42 early-course, minimally medicated patients with a psychotic disorder (EC), 42 familial high risk first-degree relatives of psychosis patients (FHR), and 57 healthy controls (HC) enrolled. Of those, 17 EC, 23 FHR and 27 HC participants met inclusion criteria and completed the study (see **Supplementary Figure 1** for enrolment). Age ranges for inclusion were as follows: EC (18-35 years), FHR (12-25) and HC (12-35). Patients were recruited from Harvard affiliated hospital outpatient programs in Boston, MA. All participants gave written informed consent. The study was approved by the institutional Human Research Committees at Beth Israel Deaconess Medical Center and Massachusetts General Hospital. Early course was defined as < 5 yrs from psychosis onset and < 1 year of lifetime exposure to antipsychotic drugs (APD). Patients were required to be unmedicated or on stable doses of antipsychotic and adjunctive medications for at least 6 weeks prior to study enrolment. Based on the Structured Clinical Interview for DSM-IV TR (SCID; First et al., 2015) or the Affective Disorders and Schizophrenia for School-Age Children-Present and Lifetime version (K-SADS-PL; Kaufman et al., 1997) and review of medical records, EC participants were characterized into schizophrenia (ECsz; n = 7) or other psychotic disorders (ECnsz; n = 10) groups (**Supplementary Table 1** for diagnosis and medication information). Diagnoses were determined by consensus between clinical assessors in diagnostic review meetings chaired by Dr. Keshavan. The familial high risk (FHR) group consisted of individuals with no personal history of psychosis (confirmed with SCID or K-SADS-PL) but a first-degree relative with a diagnosis of schizophrenia (n = 20) or schizoaffective disorder (n = 3) based on interview of either the affected relative or a reliable informant other than the index subject by FHR experienced psychiatrist (MSK), and supplemented by medical records, when available. FHR participants were confirmed to not be taking any antipsychotic medications. Healthy controls (HC) were recruited from the community by advertisements and were excluded for a personal history of mental illness confirmed by the SCID-Non-Patient Edition, a family history of psychotic disorders, and current treatment with any medication known to affect sleep or cognition.

General exclusion criteria included substance abuse or dependence within the past 6 months; unstable medical conditions that may affect sleep (e.g., diabetes, thyroid disease); pregnancy/breast feeding; a history of head injury resulting in prolonged loss of consciousness (> one day) or neurological sequelae; intellectual disability; a diagnosed sleep disorder other than insomnia; neurological disorders such as epilepsy. All participants also completed a screening test for the finger tapping motor sequence task (MST), see below. This screening test required participants to type at least 24 correct sequences of “1-2-3-4” during two 30 second trials with the left hand to be eligible for the main study. Healthy controls were demographically matched to the EC and FHR groups on age (*F* (2, 64) = 1.63, *p* = .20) and sex (χ^2^ (2) = 3.56, *p* = .17). There were significant group differences in subjective sleep quality (*F* (2, 63) = 9.35, *p* < .001), with both EC and FHR participants reporting subjectively worse sleep quality than HC (*p*s < .005). ECsz and ECnsz were matched in terms of age (*t* (17) = 1.47, *p* = .16) but not mean parental education (*t* (17) = 3.48, *p* = .004; lower in ECsz) or the severity of positive or negative symptoms (all *Ps* < .007; more severe symptoms in ECsz). Demographics and clinical characteristics are shown in **Table 1**.

**Table 1.** Sample demographics and clinical characterization.

|  | HC | FHR | EC | stat | *p* |
| --- | --- | --- | --- | --- | --- |
| Age (yrs) | 24 (4) | 23 (6) | 21 (4) | 1.63 | .20 |
| Sex (%) |  |  |  |  |  |
| Female | 47.1 | 69.6 | 44.4 | 3.56 | .16 |
| Male | 52.9 | 30.4 | 55.6 |  |  |
| Ethnicity (%) |  |  |  |  |  |
| Hispanic or Latino | 18.5 | 4.3 | 0 | 5.30 | .07 |
| Not Hispanic or Latino | 81.5 | 95.7 | 100 |  |  |
| Race (%) |  |  |  |  |  |
| Asian | 18.5 | 17.4 | 5.9 |  |  |
| Black or African American | 7.4 | 13.0 | 41.2 | 11.77 | .07 |
| White | 63.0 | 69.6 | 47.1 |  |  |
| Other, not specified | 11.1 | 0 | 5.8 |  |  |
| Marital status (%) |  |  |  |  |  |
| Married or sustained relationship | 7.4 | 13.0 | 5.9 | 0.75 | .69 |
| Never married | 92.6 | 87.0 | 94.1 |  |  |
| Mean parental education (yrs) | 16.0 (2.75) | 15.9 (3.44) | 14.8 (3.3) | 0.79 | .46 |
| Handedness <sup>a</sup> | 60.6 (50.2) | 49.5 (62.7) | 55 (54.4) | 0.23 | .80 |
| Subjective sleep quality (PSQI <sup>b</sup>) | 2.63 (1.98) | 5.64 (4.23) | 7.12 (4.4) | 9.34 | < .001 |
| Diurnal preference (MEQ <sup>c</sup>) | 51.1 (11.4) | 47.9 (10.2) | 48 (10.7) | 0.67 | .52 |
| Symptom Severity (PANSS <sup>d</sup>) |  |  |  |  |  |
| PANSS total | - | 36.0 (6.14) | 53.1 (15.4) | 22.84 | < .001 |
| PANSS positive | - | 7.27 (0.63) | 11.6 (4.91) | 17.22 | < .001 |
| PANSS negative | - | 8.77 (2.39) | 14.7 (6.01) | 17.91 | < .001 |
| PANSS general | - | 19.9 (4.45) | 26.8 (6.76) | 14.54 | < .001 |
| Psychosis proneness (Chapman) |  |  |  |  |  |
| Total | 10.2 (9.7) | 12.9 (8.22) | 34 (21.4) | 18.57 | < .001 |
| Perceptual aberration | 1.22 (2.7) | 2.74 (4.42) | 8.82 (9.05) | 10.47 | < .001 |
| Magical ideation | 2.19 (3.5) | 3.09 (1.88) | 11.5 (7.22) | 26.40 | < .001 |
| Social anhedonia | 6.78 (5.6) | 7.09 (5.28) | 13.7 (9.16) | 6.59 | .003 |
| Estimated IQ |  |  |  |  |  |
| WASI-II | 106 (13.1) | 112 (15.02) | 101 (13.3) | 2.54 | .09 |
| AMMONS | 100 (11.2) | 106 (15.1) | 97.2 (12.6) | 2.16 | .12 |

Note. Sex, ethnicity, race, and marital status are shown as a percentage of the sample. All other data are displayed as mean (standard deviation). <sup>a</sup>Handedness calculated with the Edinburgh handedness inventory (Oldfield, 1971), where a higher number indicates stronger right-hand preference (theoretical range -100 - 100). <sup>b</sup>Total score on the Pittsburgh sleep quality index (PSQI, Buysse et al., 1989). Higher score indicates worse subjective sleep (theoretical range 0-21). <sup>c</sup>Morningness-Eveningness Questionnaire (MEQ, Horne & Ostberg, 1976). Higher score indicates stronger morning preference (theoretical range 16-86). <sup>d</sup>Positive and Negative Syndrome Scale (PANSS, Higher score indicates more severe symptoms (total scale theoretical range 30-210). Chapman scale. Higher score indicates higher endorsement of psychotic-like symptoms (total scale theoretical range 0-104) WASI-II = Wechsler Abbreviated Scale of Intelligence-Revised. AMMONS = Ammons quick test. Group differences assessed via ANOVA or chi-square test as appropriate. Patients showed significantly higher symptomatology on total PANSS and all subscales than relatives. On all Chapman scales, the main effect of group was driven by higher symptomatology in patients compared to both relatives and controls ( $ps < .004$ ), with no significant difference between relatives and controls ( $ps > .33$ ).

### Procedures

#### Study overview

All participants completed an initial screening visit to provide informed consent, diagnostic interviewing, clinical characterization (symptom ratings obtained from the positive and negative symptom scale, PANSS; Kay et al., 1987) and perform the MST screening test. If they met criteria for inclusion, they returned for cognitive assessments. IQ was estimated with the Wechsler Abbreviated Scale of Intelligence (WASI-II) (Wechsler, 2011) and the AMMONS IQ test (Ammons & Ammons, 1962). All participants also completed a battery of self-report questionnaires, including the Chapman Psychosis Proneness Scale (Chapman et al., 1976). After completing all screening and baseline assessments, participants were given a tour of the Clinical Research Center (CRC) sleep monitoring rooms to familiarize them with the overnight stay portion of the study.

Approximately 1 week after the initial visit, participants completed two experimental visits, separated by 1 week. For each visit, participants spent two consecutive nights at the CRC for polysomnography (PSG) monitoring of sleep. Prior to the visit, participants were instructed to: 1) keep their typical daily schedules and obtain adequate sleep in the week preceding the study visits and not to nap during study days (confirmed with wrist actigraphy); 2) avoid alcohol and recreational drugs 24 hours prior to visits and; 3) avoid caffeine intake on the day of the visits. The first night of each visit served as a baseline and the second as an experimental night during which participants trained on the procedural (MST) or declarative (word pairs test) memory tasks prior to bed and were tested after awakening the following morning. The MST visit always occurred first. We prioritized collection of MST data since it is the most extensively validated measure of sleep-dependent memory consolidation and chronic patients show deficits (Manoach et al., 2004, 2010; Wamsley et al., 2012). Participants engaged in their usual activities during the day. **Figure 1** shows an overview of the study protocol.

**Figure 1.**
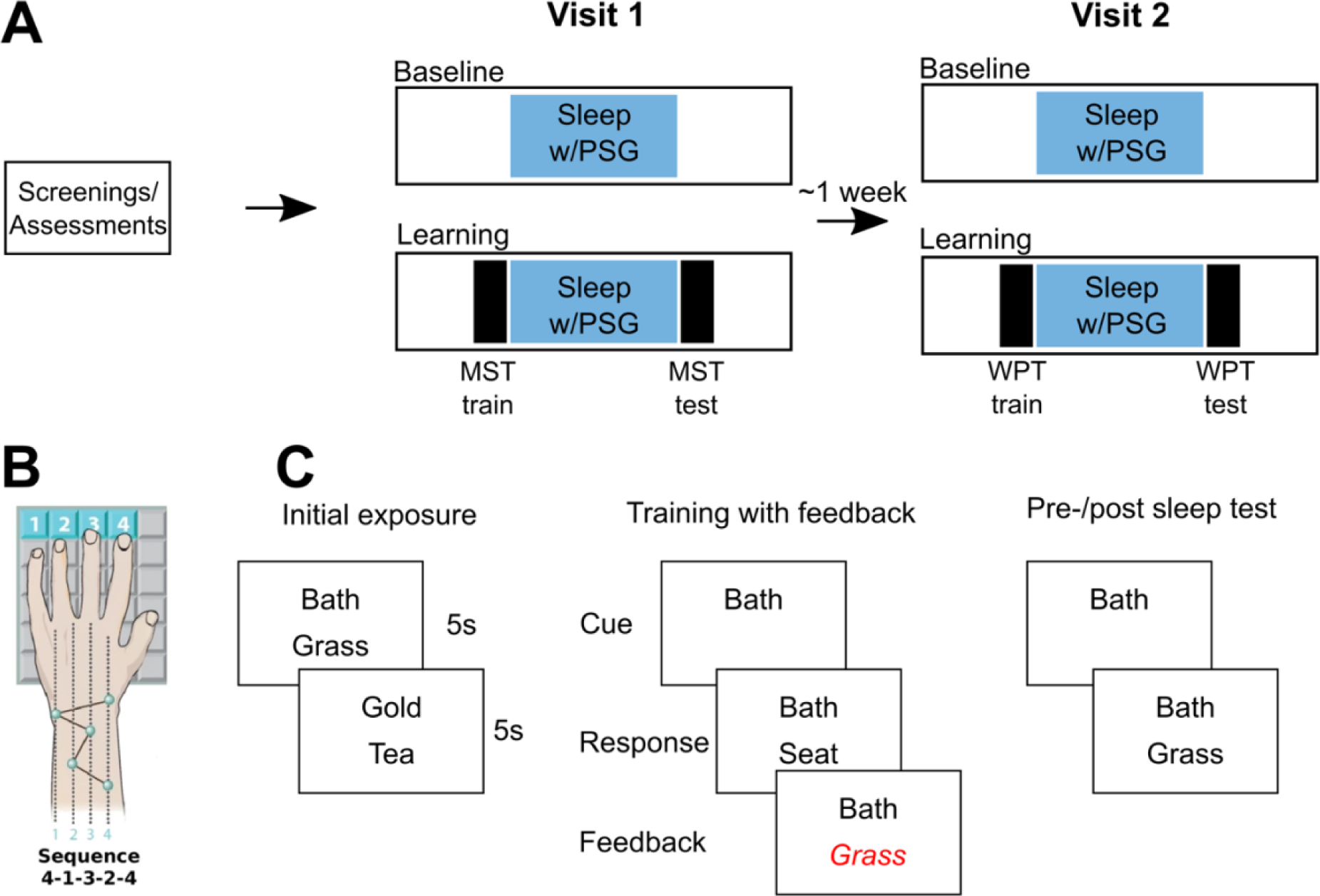
Study protocol. **A** - Timeline of the study. Following baseline screenings and assessments, which included an introductory visit to the sleep laboratory, participants took part in a four night overnight sleep protocol divided into two separate visits approximately one week apart. Each visit consisted of two consecutive nights, a baseline night and a learning night. On Visit 1, the learning night consisted of training on the motor sequence task (MST) prior to sleep, and being tested the following morning. Following post-sleep testing, participants trained and tested on a new “control” sequence. During Visit 2, the learning night consisted of training and testing on the word pairs task (WPT) in the evening, and being re-tested the following morning. **B** - Motor sequence task. In the evening participants were trained (12 trials) to type a five-digit sequence as quickly and accurately as possible. The following morning, participants were tested on an additional 12 trials after sleep. The control sequence followed the same protocol except a new five-digit sequence was used. **C** - Word pairs task. Participants were exposed to pairs of words. Following initial exposure, participants were shown the top word of each pair and were asked to recall its associate. If they answered incorrectly, participants were shown the correct word. This procedure continued until participants correctly recalled 60% of the target words. Before and after sleep, participants performed a cued recall test where each cue word was shown once, and participants had to recall the target word. No feedback was given on these tests.

#### Motor sequence task (MST)

The MST requires participants to press four numerically labelled keys with the fingers of their left hand, repeating a five-element tapping sequence (e.g. 4-1-3-2-4) as quickly and as accurately as possible for a period of thirty seconds (Walker et al., 2002). To minimize working memory load, the numeric sequence was displayed in white against a green background at the top of the screen. The time and value of each key press was recorded, and a dot was displayed on the screen for each key press. Each 30 second trial was automatically scored for the number of correct sequences and the number of errors. Each tapping trial was followed by a 30 second rest period, during which the screen was red and the number of seconds remaining before the start of the next trial was displayed as words. After the countdown reached three, the words were replaced with tones, one per second, leading to the start of the next trial. Each session consisted of 12 trials, with one session taking place in the evening before sleep (Training Session), and a second identical session taking place the following morning (Test Session). Performance was measured as the number of correctly typed sequences per trial. To assess for differences in practice-dependent learning during the evening training session, we calculated the degree of learning during training as the percent increase in correctly typed sequences from the first training trial to the average of the last three included trials of training. To examine sleep-dependent improvement, we calculated the overnight improvement as the percent increase in correctly typed sequences from the average of the last three training trials at night to the first three test trials the following morning (Walker et al., 2002).

#### Word pair task (WPT)

Stimuli consisted of single-syllable, high-frequency, concrete nouns that were paired to create a list of 24 semantically unrelated cue-target word pairs (Baran et al., 2016). The task consisted of four phases: initial exposure, training with feedback, immediate pre-sleep test, and delayed post-sleep test. During initial exposure, each pair was displayed on the screen for five seconds, one word above the other. During exposure, participants were instructed to remember which words go together. To avoid primacy and recency effects, only the middle 20 pairs were randomized during subsequent phases. During training with feedback, participants were shown the top word of each pair, and asked to verbally report its associate. The experimenter typed their response. If it was correct, the word was displayed on the screen below the cue word for two seconds. If the response was incorrect, the correct word was instead displayed on the screen for two seconds. This procedure was repeated until participants reached a criterion of 60% accuracy (i.e. correctly recalled at least of 12 of the middle 20 pairs), or until 15 training rounds had elapsed. The pre-sleep test immediately followed training. This phase followed the same procedure as training, except the cue word of each pair was only shown once, and no feedback was given. A second post-sleep test, following the same procedure as the pre-sleep test, was administered the following morning. Outcome measures were: Number of rounds taken to reach criterion performance, recall performance at the immediate test, and overnight change in memory. Overnight change in word pair memory was calculated as the relative change in recall between delayed and immediate tests [(delayed recall - immediate recall) / immediate recall)].

### Polysomnography

Data were acquired at 400 Hz with either an Aura LTM64 (Grass Technologies, Astro-Med, Inc., RI) or a Natus Embla RDx (Natus Medical Inc, CA) system and EEG caps (Easycap GmbH, Herrsching, Germany) with 58 EEG electrodes positioned in accordance with the 10-20 system. Additionally, electrodes were placed on the left and right mastoids, above the right eye and below the left eye (for EOG measures), two placed on the chin (for EMG measurements), one on the forehead (recording reference), and one on the collarbone (ground). All impedances were kept < 25 kOhm. Sleep scoring was performed to standard criteria, with each 30 second epoch being visually scored by a trained expert as Wake, N1, N2, N3 or REM. Scorers were blind to group, visit, and condition. Artifacts were identified and removed automatically using Luna (https://zzz.bwh.harvard.edu/luna/). EEG data were preprocessed using custom MATLAB scripts and EEGLAB routines. Data were re-referenced to the average of the two mastoids, bandpass filtered between 0.3-35Hz, and notch filtered at 60Hz.

### Power spectral density

Estimates of power spectral density (PSD) were obtained for stage 2 NREM (N2) sleep at all electrodes. PSD was estimated using Welch’s method with 5s Hamming windows and 50% overlap. PSD estimates were derived from the derivative of the EEG time series to minimize 1/*f* scaling (Cox et al., 2017).

### Sleep spindles

Sleep spindles were automatically detected using a wavelet-based detector, validated for use in both healthy individuals and people with schizophrenia (Wamsley et al., 2012; Warby et al., 2014). The raw EEG signal was subjected to a time-frequency decomposition using complex Morlet wavelets. The peak frequency of the wavelet was set at 13.5Hz, with a 3Hz bandwidth centered on the wavelet peak. Therefore, spindles in the 12-15 Hz range were detected. This frequency range corresponds primarily to “fast” spindles. Spindles were detected on each channel by applying a thresholding algorithm to the extracted wavelet scale. A spindle was detected whenever the wavelet signal exceeded a threshold of nine times the median signal amplitude of all artifact-free data for a minimum of 400ms. The threshold of nine times the median empirically maximized between-class (“spindle” vs “non-spindle”) variance in previous samples of schizophrenia and control patients with 12-15 Hz spindles (Wamsley et al., 2012). Spindles were detected during both N2 and N3 sleep. As in prior studies, we focused on N2 sleep (Mylonas et al., 2020; Wamsley et al., 2012, 2013). Our primary outcome measure was N2 spindle density, calculated as spindles per minute. We also extracted N2 spindle amplitude, defined as the maximal voltage of a 4 sec window centered on the peak of the detected spindle.

### Slow oscillations and their coupling with spindles

Slow oscillations (SOs) were detected at every electrode during N2 and N3 sleep using an automated algorithm (Helfrich et al., 2018; Staresina et al., 2015). First, data were bandpass filtered between 0.5 - 4Hz, and all positive-to-negative zero crossings were identified. Candidate slow oscillations were marked if two such consecutive zero crossings fell 0.8 - 2 seconds apart, corresponding to 0.5 - 1.25 Hz. Peak-to-peak amplitudes for all candidate oscillations were determined, and oscillations in the top quartile (i.e. with the highest amplitudes) at each electrode were retained as slow oscillations. We extracted N2 slow oscillation density (number of slow oscillations per minute of artifact free N2 sleep), as well as slow oscillation peak-to-peak amplitude (µV).

To identify SO-spindle coupling events, EEG data were bandpass filtered, once between 0.5 - 4 Hz and once between 12 - 15 Hz. Then, the Hilbert transform was applied to extract the instantaneous phase of the delta (0.5 - 4 Hz) filtered signal and the instantaneous amplitude of the sigma (12 - 15 Hz) filtered signal. For each detected spindle, the time of peak amplitude was determined. If the spindle peak occurred within the time course (i.e., between two positive-to-negative zero crossings) of a detected SO, a SO-spindle coupling event was scored, and the phase angle of the slow oscillation at the peak of the spindle was determined. Finally, we extracted the number, density and percentage of coupled spindles, average coupling phase (in degrees) and coupling consistency, measured as the mean length of the vector plotted in polar coordinates (Berens, 2009). As with sleep spindles, we primarily focused on SO and coupling events that occurred during N2 sleep.

### Statistical analysis

#### Group differences on sleep parameters

Linear mixed-effects models were run to investigate group differences in sleep parameters at each electrode with Group (HC, FHR, EC), Visit (Visit 1, Visit 2), Night (Baseline, Learning), and their interactions entered the model as fixed effects and participant entered as a random effect. In all models, age was included as a continuous covariate.

To assess differences in coupling phase, circular two-way ANOVA models were used to first assess the effect of Group and Visit (Baseline Visit 1, Baseline Visit 2) and their interaction on coupling phase. To elucidate potential differences in Night, further circular ANOVA models were implemented with Group and Session (Baseline, Learning) and their interaction as factors, performed separately for Visit 1 and Visit 2.

#### Group differences in sleep-dependent memory consolidation

MST: As an initial pre-processing step, outlier MST trials were removed. Outlier trials were detected for each participant, by first fitting a piecewise exponential model to each participant’s learning curve (Mylonas et al., 2020). Specifically, each participant’s MST data were fit to the equation:

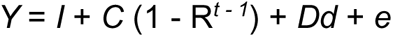

Where *Y* = correct sequences typed on trial *t; I* = initial performance (average score on trial 1); *C* = change in performance from trial 1 to the asymptote (amount learned); 1 - *R* = the learning rate; *t* = trial number; *D* = overnight improvement; *d* = 0 (for evening) or *d* = 1 (for morning); and *e* is a stochastic error term. The squared residuals for each trial were calculated, and trials falling more than 2 standard deviations below the average of the distribution of the squared residuals across all trials were excluded. Only data points below the estimated learning curve were excluded, since we did not have grounds to exclude trials where participants performed better than expected. One-way ANCOVA was used to assess group differences in practice-dependent learning and sleep-dependent improvement with the factor Group (HC, FHR, EC) and age entered as a covariate.

WPT: Group differences in number of rounds taken to reach criterion performance, recall performance at the immediate test, and overnight change in memory performance were assessed via a series of one-way ANCOVAs with the factor Group (HC, FHR, EC), and age entered as a covariate.

#### Associations between sleep spindle density and overnight memory consolidation

We used spindle measures from the learning nights to examine relationships between sleep and overnight consolidation. In separate models, we regressed MST improvement and WPT change in recall against spindle density at each electrode, with Group included as a fixed effect and age added as a continuous covariate. Robust linear regression procedures were used to minimize the influence of outliers. To determine whether spindles coupled to slow oscillations were a better predictor than uncoupled sleep spindles, the goodness of fit of two models was compared (Mylonas et al., 2020). In one model, memory consolidation was predicted from uncoupled spindle density, and in a second model memory was predicted from coupled spindle density. The goodness of fit of each model at each electrode was compared using the Bayesian Information Criterion (BIC), where a lower value indicates a better model fit (Schwarz, 1978).

#### Associations between sleep spindle density and symptomatology

Relationships between sleep spindles and measures of symptomatology (PANSS scores, FHR and EC only, Chapman scales all participants) were assessed using the same robust linear regression procedure described above.

#### Multiple comparisons correction

To control for multiple comparisons across electrodes, and to take into account the spatial correlation of the EEG data, we used a cluster-based permutation method (Maris & Oostenveld, 2007). For each term in the model, clusters were formed from adjacent electrodes that met an uncorrected threshold of *p* < .05. Permutation distributions were created by randomly shuffling the labels (Group, Visit, Session, Age) 1000 times at each electrode and retaining the cluster with the maximum statistic for each permutation (Mylonas et al., 2020). The cluster-level corrected *p*-value is the probability that the observed cluster would be found by chance within the permutation distribution. A cluster-corrected *p* < .05 was deemed statistically significant. Follow-up pairwise estimated marginal means tests were used as appropriate to further delineate significant group differences. Pairwise tests were performed on data averaged across significant electrodes in the overall cluster, with age included as a covariate. To examine whether any observed deficits in EC were driven by schizophrenia patients, pairwise estimated marginal means tests were performed between ECsz and ECnsz participants, again using data averaged across electrodes in the overall cluster.

## Results

All results remained unchanged in models that did not include age as a covariate. Similarly, all group differences in N2 sleep physiology replicated when analyses were run on N3 sleep (reported in supplement), and when the three FHR participants whose relative had a non-schizophrenia psychosis were removed.

### Sleep quality and architecture

There were no significant group differences in sleep quality or architecture (Table 2 displays statistics averaged across all nights; see **Supplementary Table 2** for each night separately. Total sleep time (TST), sleep onset latency, sleep efficiency, and wake after sleep onset did not significantly differ among groups (all *p’*s > .13). A borderline main effect of group was seen for time spent in REM sleep (*p* = .051), with lower REM time in FHR and EC compared to HC (**Table 2**). No other main effects of group were seen for time or percentage of TST spent in any other sleep stage.

**Table 2.** Sleep architecture.

|  | HC | FHR | EC | *F* | *p* |
| --- | --- | --- | --- | --- | --- |
| Total sleep time (min) | 508 (74) | 486 (67) | 507 (54) | 1.79 | .18 |
| Sleep onset latency (min) | 25 (29) | 33 (36) | 20 (20) | 1.18 | .32 |
| Sleep efficiency (%) | 86 (8) | 83 (11) | 85 (9) | 2.06 | .14 |
| Wake after sleep onset (min) | 52 (41) | 57 (48) | 65 (49) | 1.72 | .19 |
| N1 (min) | 31 (23) | 28 (23) | 34 (40) | 0.18 | .84 |
| N2 (min) | 262 (54) | 251 (63) | 259 (57) | 1.34 | .27 |
| N3 (min) | 102 (31) | 111 (46) | 121 (47) | 1.66 | .20 |
| REM (min) | 113 (33) | 96 (36) | 93 (42) | 3.17 | .05 |
| N1 (% of TST) | 6.1 (4.7) | 5.7 (4.3) | 6.9 (8.2) | 0.14 | .87 |
| N2 (% of TST) | 51.5 (6.7) | 51.6 (8.5) | 51.1 (10) | 0.30 | .74 |
| N3 (% of TST) | 20.5 (6.5) | 23.4 (9.2) | 23.8 (8.6) | 2.52 | .09 |
| REM (% of TST) | 21.9 (5.1) | 19.3 (5.8) | 18.1 (7.6) | 3.01 | .06 |

*Note.* Sleep architecture averaged over the four experimental nights. *F* statistic for the main effect of group on sleep architecture. For the borderline main effect of REM (min), effects were driven by reduced REM time in FHR and EC relative to HC ( $p$ 's < .001). The same trend emerged for REM (% of TST); $p$ 's < .005).

### Power spectral density

The N2 power spectrum, averaged across nights and electrodes, is displayed in **Figure 2A**. Topographical differences between groups in canonical frequency bands are shown in **Figure 2B**. For PSD in the sigma band (12-15Hz, corresponding to sleep spindle frequency), there was a significant main effect of group at 37 electrodes (*F*_sum_ = 153.26, *p* = .007), with reduced spindle-band power in EC (all *p’*s <= .004). There was no difference between ECsz and ECnsz patients (*p* = .97). No other group differences emerged in any other canonical frequency band (**Figure 2B**), demonstrating a selective spindle-band deficit in EC.

**Figure 2.**
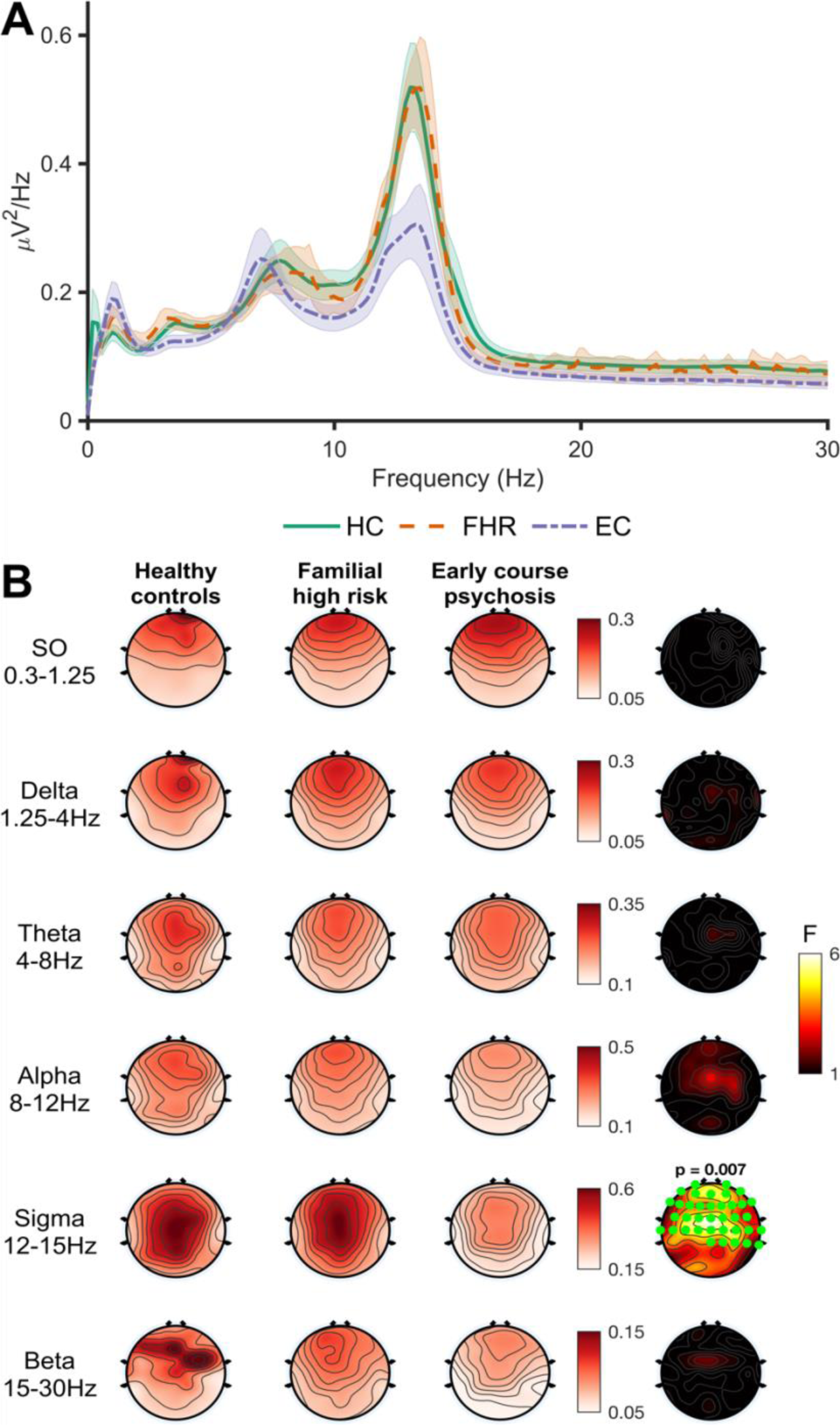
N2 sleep spectral power. **A** - Global power spectral density (PSD) in frequencies from 0-30Hz, collapsed across electrodes and nights. Shaded area around lines indicates the between-subject standard error. **B** - Main effect of group on PSD in canonical frequency bands. Topographies averaged across all four nights are shown for each group separately. Right hand topoplots show F values at each electrode for the main effect of group. Significant electrodes (cluster-corrected) are highlighted in green. Significant cluster *p* values displayed above plot.

### Sleep spindles

There was a significant main effect of group on spindle density in a cluster of 52 of the 58 electrodes (*F_sum_* = 249.12, *p* = .009; **Figure 3A, upper row**). EC exhibited a significantly lower spindle density than both HC (*t* (63) = 3.02, *p* = .004) and FHR (*t* (63) = 3.08, *p* = .003; **Figure 3B**). There was no evidence of a spindle deficit in FHR compared to HC (*t* (63) = 0.13, *p* = .90). No other effect was significant (**Supplementary Table 3**). Within EC, there was significantly lower spindle density in ECsz compared with ECnsz (*t* (14) = 2.39 *p* = .03; **Figure 3B**) who did not differ from HC (*p* = .17).

**Figure 3.**
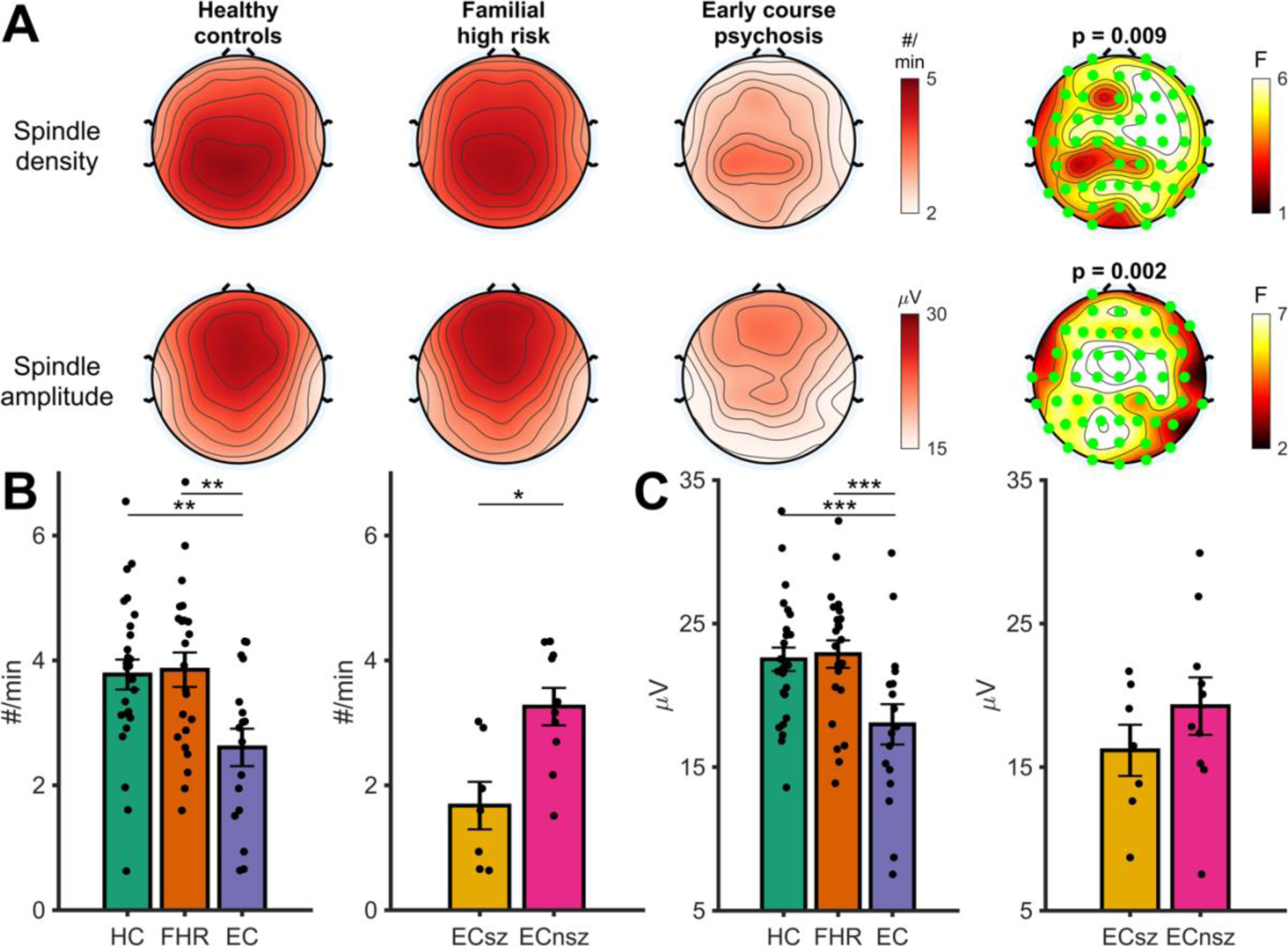
N2 sleep spindles. **A** - Topographies showing the main effect of group on N2 spindle density (Row 1) and amplitude (Row 2). N2 spindle topographies (averaged across all four nights) are shown for the three groups separately. The right-hand plot shows F values at each electrode for the main effect of group. Significant electrodes (cluster-corrected) are highlighted in green. Cluster *p-*value displayed above plot. **B** - Pairwise tests for the main effect of group on N2 spindle density with spindle density averaged over significant electrodes in the cluster highlighted in A. Left hand plot shows differences between the three groups. Right hand plot shows differences between schizophrenia and non-schizophrenia psychosis patients. **C** - Same as B, but for N2 spindle amplitude. HC = Healthy controls, FHR = First degree relatives, EC = Early course psychosis, ECsz = Early course schizophrenia, ECnsz = Early course non-schizophrenia psychosis. Error bars indicate the standard error. *** = *p* < .001, ** = *p* < .01, * = *p* < .05 from pairwise estimated marginal means tests.

A significant main effect of group on spindle amplitude was observed in a cluster of 53 electrodes (*F_sum_* = 309.80, *p* = .002; **Figure 3A, lower row**) with significantly reduced spindle amplitude in EC (**Figure 3C**) compared to both HC (*t* (63) = 3.49, *p<* .001) and FHR (*t* (63) = 3.51, *p* < .001). As with spindle density, spindle amplitude did not differ between HC and FHR (*t* (63) = 0.08, *p* = .94). There were no significant interactions between group and either visit or session (**Supplementary Table 3**). There was no difference in spindle amplitude between ECsz and ECnsz (t (14) = 1.34, p = .21; **Figure 3C**).

To summarize, EC showed both reduced density and amplitude of N2 spindles compared with HC and FHR, who did not differ in spindle density or amplitude. In the case of spindle density, the group differences were driven by ECsz.

### Slow oscillations and SO-spindle coupling

There were no group differences in either the density or peak-to-peak amplitude of detected SOs (**Figure 4A**; **Supplementary Table 3**). We next examined the properties of SO-spindle coupling. Of all detected spindles, 11.6% ± 5.4% were coupled to an SO. Across participants and at all electrodes, we found significant non-uniformity in the preferred phase of the SO to which spindles were coupled (all Z >= 8.65, all p < .001). Sleep spindles preferentially coupled to the rising phase of the SO, close to the positive peak (M = -28.7° ± 19.7°) at electrode Cz; **Figure 4B)**. There were no group differences in the average coupling phase of sleep spindles to SOs (**Supplementary Table 3**) or with the number, density, or the percentage of spindles coupled to SOs at any of the electrodes (**Supplementary Table 3**).

**Figure 4.**
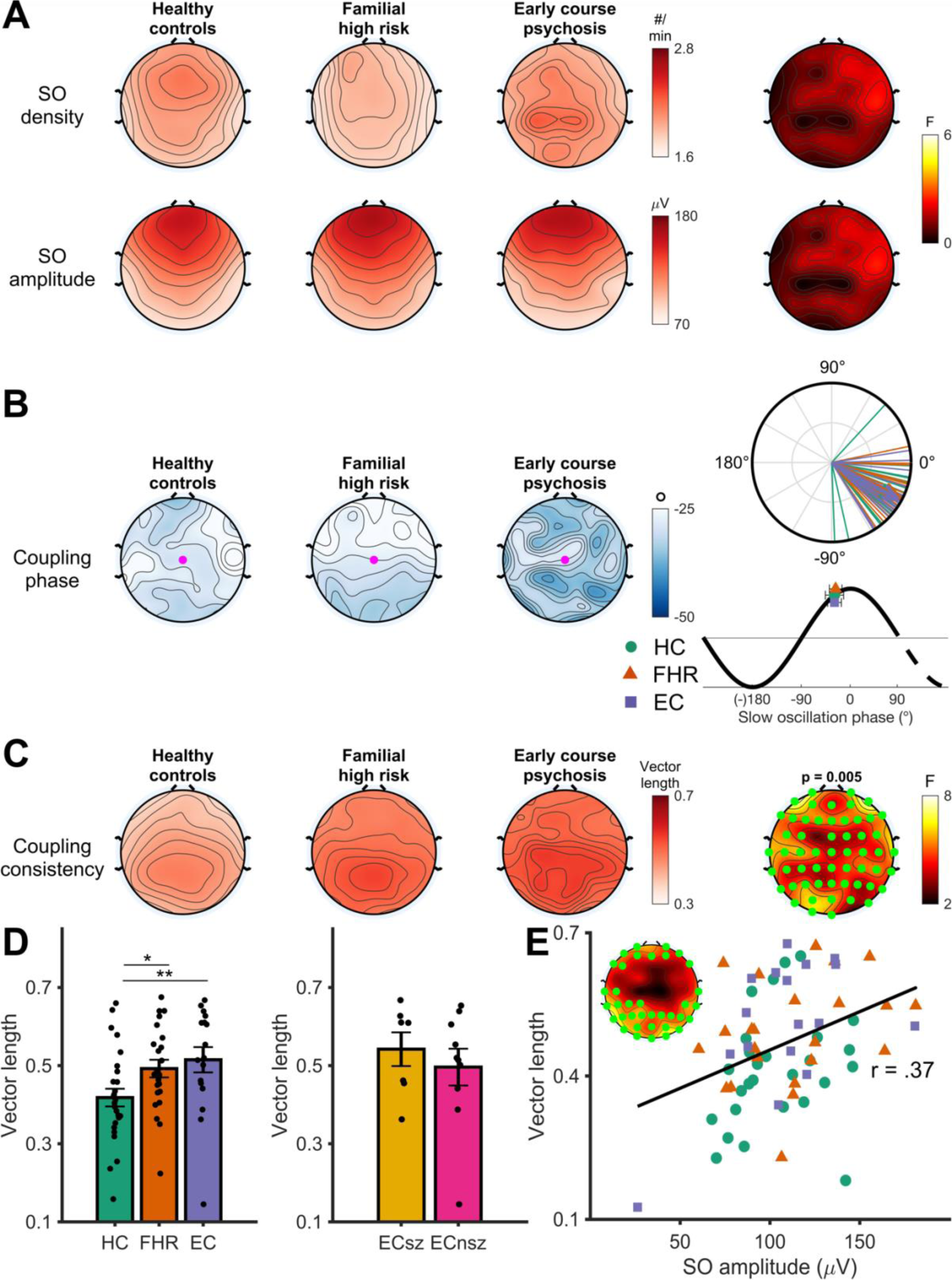
N2 slow oscillation spindle coupling. **A** - Slow oscillation density (top row) and peak-to-peak amplitude (bottom row) in each group, averaged across the four nights. Rightmost topoplots display F values for the main effect of group. **B** - Preferred coupling phase of spindles to slow oscillations (SOs) in each group, averaged across the four nights. Right: circular phase plot displaying phase distributions across participants for each group at electrode Cz (highlighted electrode in topography). Each line indicates the preferred coupling phase of individual participants. The direction of the arrow indicates the average phase across participants, separately for each group, with the length of the arrow indicating the coupling consistency across participants. A coupling of phase of 0° indicates preferential coupling of spindles at the positive peak of the slow oscillation. A coupling phase of 180° indicates preferential spindle coupling at the negative trough of the slow oscillation. Mapping of SO phase to topographical and circular plots is below the circular phase plot. **C** - Topographies showing the main effect of group on coupling consistency. Coupling consistency (measured as the mean vector length) topographies (averaged across all four nights) are shown for the three groups separately. The right hand plot shows F values at each electrode for the main effect of group. Significant electrodes (cluster-corrected) are highlighted in green. Cluster *p* value displayed above plot. **D** - Pairwise tests for the main effect of group on coupling consistency, with coupling consistency averaged over significant electrodes in cluster.. HC = Healthy controls, FHR = First degree relatives, EC = Early course psychosis, ECsz = Early course schizophrenia, ECnsz = Early course non-schizophrenia psychosis. Error bars indicate the standard error. ** = *p* < .01, * = *p* < .05 from pairwise estimated marginal means tests. **E** – Robust linear regression between slow oscillation amplitude and spindle coupling consistency. Insert shows significant electrodes.

There was a significant group effect on coupling consistency in a cluster of 56 electrodes (*F_sum_* = 277.61, p = .005; **Figure 4C**; **Supplementary Table 3**) corresponding to significantly *lower* SO-spindle coupling consistency in HC compared with both EC (t (63) = 2.98, p = .004) and FHR (t (63) = 2.37, p = .021; **Figure 4D**). There were no differences between EC and FHR (t (63) = 084, p = .40), nor were there significant differences between ECsz and ECnsz (t (14) = 1.17, *p* = .26). Across all participants, the consistency of SO-spindle timing correlated with slow oscillation amplitude, with a higher slow oscillation amplitude predicting more consistent coupling (r = 0.37, p = .002; **Figure 4E**).

Except for the greater consistency of the SO phase at which spindles peaked in both EC and FHR groups, EC and FHR groups did not differ from controls in spindle-SO coupling.

### Overnight memory consolidation

#### MST

MST learning curves are displayed in **Figure 5A-B**. Although EC typed fewer correct sequences overall, there were no group differences in improvement during training (*F* (2, 63) = 1.54, *p* = .22), including between ECsz and ECnsz patients (*F* (1, 14) = 0.14, *p* = .79). For overnight improvement, there was no overall main effect of group (*F* (2, 63) = 0.75, *p* = .48). However, a direct contrast between ECsz and all other groups revealed significantly less overnight improvement in ECsz (*t* (64) = 2.17, *p* = .034; **Figure 5B-C**). In ECsz, the degree of overnight improvement (6.3% ± 16.6%) was not significantly different from zero (*t* (6) = 1.01, *p* = .35), whereas all other groups showed substantial improvement (18.2% ± 13.5%; *t* (60) = 10.4, *p* < .001).

**Figure 5.**
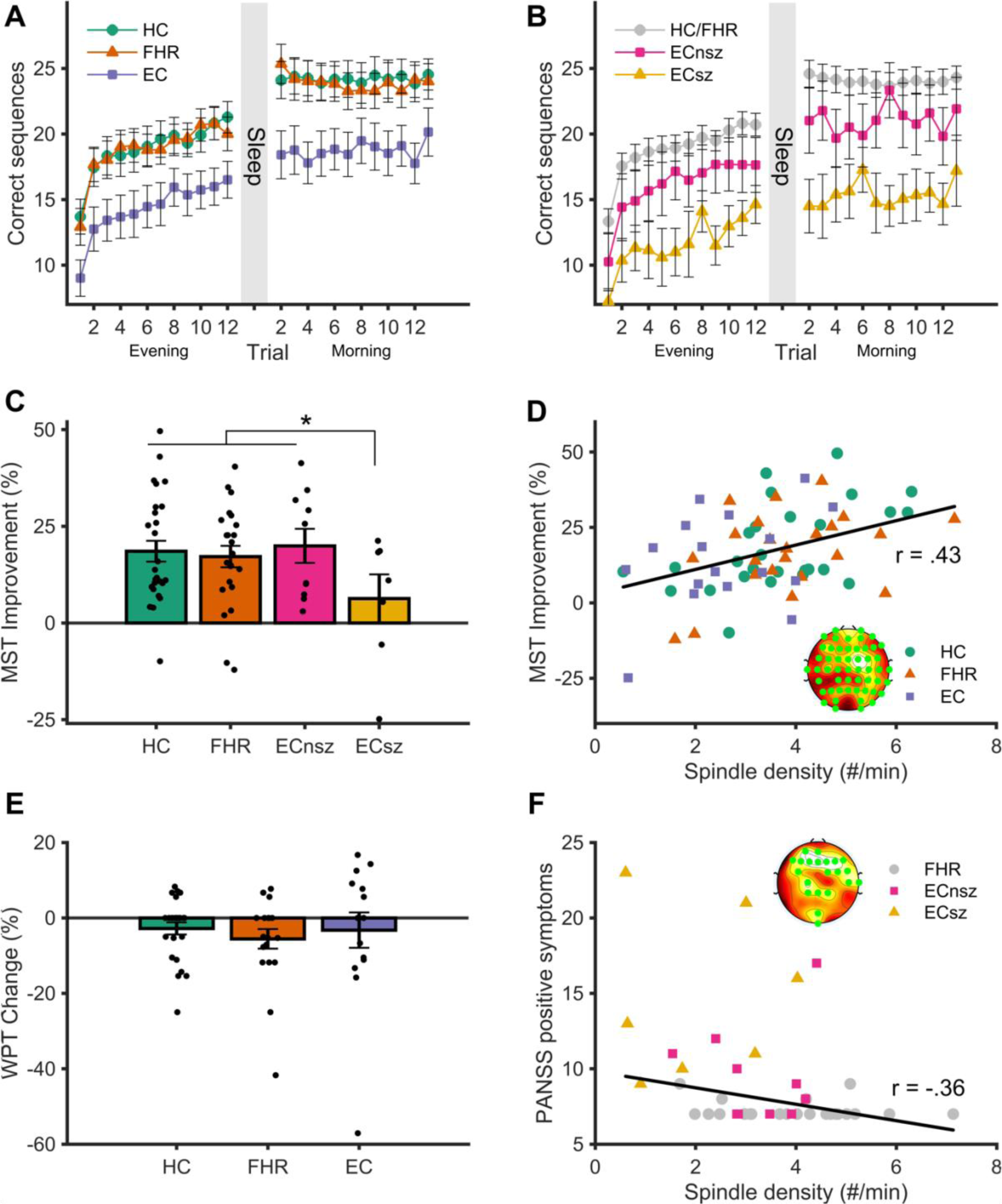
Overnight memory consolidation and spindle correlations. **A** - Motor sequence learning across training (evening) and test (morning) trials for each group. The Y axis represents the number of correctly typed sequences per trial. **B** - Motor sequence learning plotted separately for schizophrenia and non-schizophrenia psychosis patients. Grey circles show the combined healthy control and familial high-risk groups. **C** - Overnight MST improvement, calculated as the percent increase in correctly typed sequences from the average of the last three training trials at night to the first three test trials the following morning. * = *p* < .05, error bars reflect the standard error. **D** – Robust linear regression between N2 spindle density and overnight MST improvement (cluster averaged). Insert shows significant electrodes in the cluster. **E** – Overnight change in word pair memory. Error bars indicate the standard error. **F** – Robust linear regression between N2 spindle density and positive symptoms as assessed by the PANSS (cluster averaged). Insert shows significant electrodes in cluster.

Higher spindle density predicted better overnight improvement (52 electrodes, *F_sum_* = 341.95, *p* = .008; **Figure 5D**). Averaging spindle density across all significant electrodes resulted in a cluster averaged correlation coefficient of *r* = .43. The strength of association between spindle density and overnight improvement did not significantly differ among groups, although the correlation was numerically stronger in HC when averaged over the significant cluster (HC: *r* = .51, FHR: *r* = .38; EC: *r* = .37; ECsz: *r* = .19, ECnsz: *r* = .32). Although coupled spindle density also correlated with MST improvement (*r* = .44, *p* < .001), coupled spindle density was not a better predictor of memory consolidation than uncoupled spindle density (*t* (57) = 1.01, *p* = .32).

#### WPT

EC took more trials to reach criterion than FHR, and numerically more than HC (main effect of group: *F* (2, 59) = 5.45, *p* = .007). However, immediate recall accuracy did not differ between the groups (*p* = .127), and there was no difference in overnight change in recall (*p* = .696; **Figure 5E**). There was no difference between ECsz and ECnsz patients in the number of trials taken to reach criterion, immediate recall accuracy, or overnight change in memory (all *p* >= .28). Neither spindle density, amplitude, nor SO-spindle coupling predicted overnight change in WPT. Exploratory analyses of N3 sleep revealed a positive correlation between N3 coupled spindle density and overnight retention of word pairs in the EC group only (*r* = .60, *p* = .022; see **Supplementary materials** for details).

### Correlations between spindles and symptomatology

Lower spindle density predicted greater positive symptom severity (PANSS) in two clusters (cluster 1: 19 electrodes, *F_sum_* = 90.18, *p* = .005; cluster 2: 2 electrodes, *F_sum_* = 8.84, *p* = .018), in the FHR and EC groups (cluster averaged correlation coefficient of *r* = -.36; **Figure 5F**). Spindle density was not associated with negative (all *p* < .16) or general PANSS scores (no cluster formed). Across all groups, spindle amplitude negatively correlated with Chapman total score (42 electrodes, *F_sum_* = 273, *p* = .007, cluster averaged correlation coefficients whole sample: *r* = -.37, HC: *r* = -.28, FHR: *r* = -.39; EC *r* = -.07; ECsz: *r* = -.52, ECnsz: *r* = .15). Of the Chapman subscales, a significant cluster only emerged for the magical ideation subscale (51 electrodes, *F_sum_* = 417, *p* = .003, cluster averaged correlation coefficients whole sample: *r* = -.41, HC: *r* = -.27; FHR: *r* = .05; EC: *r* = -.30; ECsz: *r* = -.60; ECnsz: *r* = -.14). No associations were found between chlorpromazine equivalent antipsychotic medication dosage and sleep spindle activity (no clusters formed, see **Table S1** for equivalent dosages).

## Discussion

We examined sleep oscillations in relation to sleep-dependent memory consolidation in early-course minimally medicated psychosis patients and young, non-psychotic first-degree relatives. Replicating previous work, and extending it to early course patients with schizophrenia, we found pronounced deficits in sleep spindle density and overnight procedural memory consolidation patients relative to controls (Demirlek & Bora, 2023; Lai et al., 2021). However, neither of these deficits extended to early course patients with other psychoses or non-psychotic first-degree relatives. Although the amount and average phase of SO-spindle coupling did not differ between the groups, both patients and relatives showed increased consistency of coupling compared to controls, replicating previous findings and extending them to early course and relative groups (Kozhemiako et al., 2022; Mylonas et al., 2020).

Sleep spindle deficits are a reliable (see meta-analysis by Lai et al., 2021) and generalizable (across racially and ethnically distinct samples; Kozhemiako et al., 2022) feature of schizophrenia. Our previous work found evidence of a spindle deficit in first-episode, antipsychotic naive patients (Manoach et al., 2014), which were able to replicate in the present sample of early course psychotic patients. For spindle density, this deficit was driven by patients with schizophrenia.

Sleep spindles are initiated in the thalamic reticular nucleus (TRN), the major inhibitory nucleus of the thalamus (Fernandez & Luthi, 2020; Guillery & Harting, 2003). A reduction of TRN-mediated inhibition of thalamocortical neurons is thought to decrease the rebound burst firing that synchronize and propagate spindle events, resulting in a spindle deficit (Clawson et al., 2016; Fernandez & Luthi, 2020). TRN abnormalities in schizophrenia have been revealed by post-mortem studies (Court et al., 1999; Smith et al., 2001; Steullet et al., 2018), and neuroimaging work has consistently identified functional hyperconnectivity between thalamus and sensorimotor cortex in patients with psychotic disorders and those at clinical high risk (Anticevic et al., 2015; Giraldo-Chica & Woodward, 2017). This hyperconnectivity may reflect a reduction of TRN inhibition of thalamocortical neurons. Indeed, we have previously reported that thalamocortical hyperconnectivity correlates with lower spindle density in schizophrenia (Baran et al., 2019). Together these findings support the hypothesis of reduced thalamic inhibition in schizophrenia during sleep and wake (Avram et al., 2018; Ferrarelli et al., 2010; Ferri et al., 2018; Manoach et al., 2014; Wamsley et al., 2012).

Spindle deficits in schizophrenia have been linked to deficits in sleep-dependent memory consolidation (Göder et al., 2015; Wamsley et al., 2012). We extended these findings to show reduced procedural memory consolidation in minimally medicated early-course schizophrenia patients, making it difficult to attribute these findings to disease chronicity. As in previous studies, the degree of overnight improvement correlated with spindle density and the density of SO-coupled spindles, consistent with a mechanistic relationship between spindles and memory consolidation deficits in schizophrenia (Manoach & Stickgold, 2019). Importantly, patients showed intact learning during training, restricting the deficit to overnight memory consolidation.

In the same sample in whom spindle deficits predicted impaired procedural memory, there were no group differences in the consolidation of a declarative word pairs task. Although deficits in verbal memory have been previously reported in schizophrenia (Baran et al., 2018; Göder et al., 2015; Weinhold et al., 2022) in the present study, consolidation of the word pairs task was not affected by the spindle deficit observed in patients. Despite this, a significant correlation of consolidation with the density of SO-coupled N3 spindles was seen in the patient group only. The fact that such associations were seen in N3 and not N2 is congruent with the finding that declarative memory consolidation is more supported by N3 sleep processes (Ackermann & Rasch, 2014).

Contrary to our hypotheses, neither sleep spindles nor memory consolidation was impaired in the first-degree relatives of patients. Compared to our previous study which reported spindle deficits in relatives (Manoach et al., 2014), in the present study the relatives were older and therefore more likely to be beyond the age of maximum risk. They may also have been healthier. However, work by other groups have observed spindle deficits in both young adult and middle aged samples of first-degree relatives of schizophrenia patients (D’Agostino et al., 2018;) including in those without any psychopathology (Schilling et al., 2017). As relatives are hypothesized to show attenuated deficits compared to probands with schizophrenia, based on presumably lower genetic risk, larger sample sizes may be necessary to evaluate group differences.

While no evidence of a spindle deficit was seen in relatives, both relatives and patients showed higher SO-spindle coupling consistency compared to controls. This finding replicates previous reports of chronic medicated patients, and extends it to early course patients and their first-degree relatives (Kozhemiako et al., 2022; Mylonas et al., 2020). In both healthy and schizophrenia samples, coupling consistency has been associated with enhanced memory consolidation, implying a higher coupling consistency to be desirable (Demanuele et al., 2016; Mikutta et al., 2019), however coupling did not predict MST consolidation in the present study. Across participants, a higher coupling consistency correlated with larger SOs, suggesting higher amplitude SOs are better able to group spindles into their excitable upstates.

The underlying mechanisms, and implications of, a higher coupling consistency in patients and their relatives remains unclear and is worthy of further study. One hypothesis is that coupling consistency reflects increased thalamocortical connectivity, as is seen in schizophrenia, specifically in sensory and motor cortices, and correlates with spindles (Baran et al, 2019). Alternatively, measures of EEG signal complexity that quantify the degree of regularity in the EEG time series have suggested reduced variability in the sleep EEG signal *in general* in schizophrenia (Keshavan et al., 2004). As such, the decreased variability in SO-spindle coupling could speak to a more global reduction in EEG variability. Mechanistic studies are needed to understand the relations of SO-spindle coupling consistency to memory and to the connectivity and synchronization of thalamocortical networks.

Strengths of this study include the multi-night experimental protocol, high-density EEG recordings, and stringent screening and recruitment procedures. However, our sample size, especially of early course patients, was relatively small, at least in part resulting from COVID-19 pandemic related constraints, and the unexpected findings need to be replicated. Second, even though our sample was minimally medicated, we cannot rule out the possibility that medication use affected our outcome measures (Krystal et al., 2008). Despite this, we did not observe a spindle density deficit in the medicated non-schizophrenia patients, nor was there a correlation between chlorpromazine equivalent APD dosage and spindle measures in the present study. A third limitation is that our memory tasks did not include a wake control condition. Although both tasks we used have been reliably shown to benefit from sleep (Berres & Erdfelder, 2021; Schmid et al., 2020), it was not possible to determine the magnitude of the sleep benefit compared to wake in the present sample. In order to reach a conclusion about whether sleep-dependent memory consolidation of word pairs in schizophrenia is diminished or not, we would need a wake control group to determine sleep specificity.

In conclusion, we found reduced sleep spindle density and a correlated deficit in overnight memory consolidation in minimally medicated, early course schizophrenia. These findings reinforce that deficits in sleep-dependent memory consolidation in schizophrenia are not due to disease chronicity or prolonged medication use. Although we did not find spindle deficits in unaffected relatives, both relatives and patients demonstrated reduced variation in the coupling between spindles and SOs. This highlights the need for future research to improving our mechanistic understanding of this metric and how it relates to the risk and development of schizophrenia. We conclude that abnormal NREM sleep oscillations are a reliable finding throughout the course of schizophrenia and are a potentially tractable target for interventions to improve memory.

## Acknowledgements

We are grateful to the staff of the Clinical Research Centers at Beth Israel Deaconess Medical Center and Massachusetts General Hospital, and to all of our participants.

## Data availability statement

The data underlying this article will be shared on reasonable request to the corresponding author.

## Supplementary results

### Correlations between N3 coupled spindles and WPT consolidation

A significant main effect was observed when this analysis was restricted to just coupled spindles during N3 sleep (14 electrodes, *F_sum_*= 71.22, *p* = .041). This main effect was superseded by a significant spindle * group interaction (14 electrodes, *F_sum_* = 61.83, *p* = .046; **Supplementary Figure 3**), reflecting that N3 coupled spindle density was positively associated with the overnight retention of word pairs in the EC group only (*r* = .66, *p* = .019; ecSZ: *r* = .89, *p* = .02; ecNSZ: *r* = .64, *p* = .09; HC: *r* = .03, *p* = .88; FHR: *r* = .25, *p* = .32). Coupled spindle density was a significantly better predictor of WPT consolidation (BIC: M = 465.78, SD = 3.46) than uncoupled spindle density (BIC: M = 467.97, SD = 3.25), *t* (57) = 7.43, *p* < .001, *d* = 0.65. There was no association between consolidation of the MST and consolidation of the WPT (*r* = .11, *p* = .39).

**Table S1.**
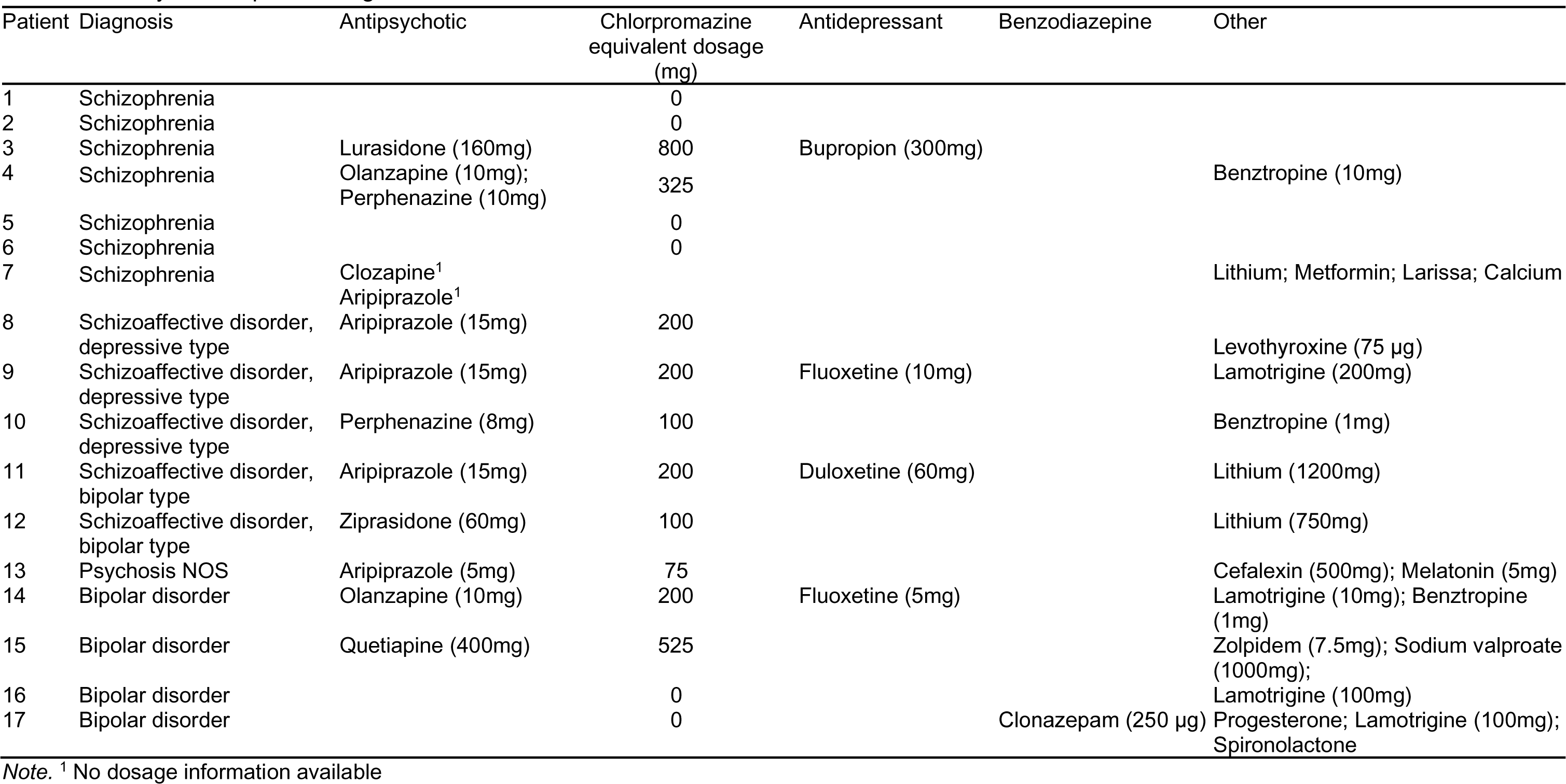
Early course patient diagnosis and medications.

**Table S2.**
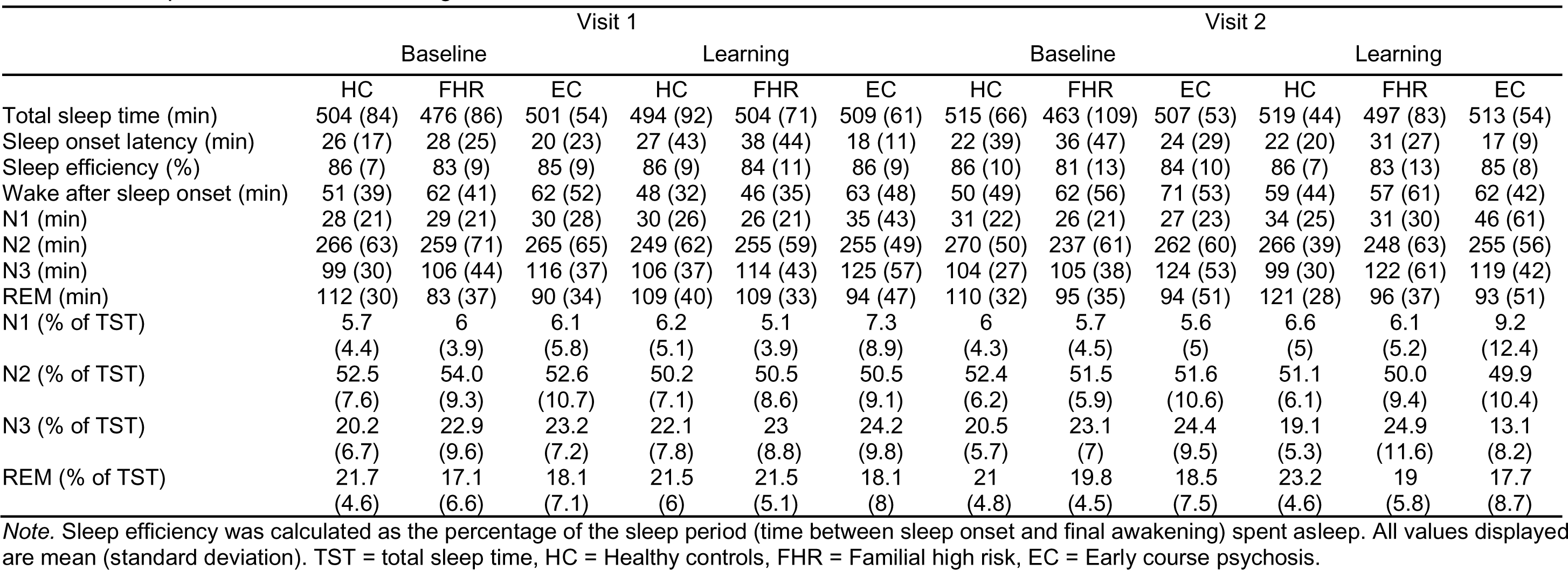
Sleep architecture for each night.

|  | Visit 1 |  |  |  |  |  | Visit 2 |  |  |  |  |  |
| --- | --- | --- | --- | --- | --- | --- | --- | --- | --- | --- | --- | --- |
|  | Baseline |  |  | Learning |  |  | Baseline |  |  | Learning |  |  |
|  | HC | FHR | EC | HC | FHR | EC | HC | FHR | EC | HC | FHR | EC |
| Total sleep time (min) | 504 (84) | 476 (86) | 501 (54) | 494 (92) | 504 (71) | 509 (61) | 515 (66) | 463 (109) | 507 (53) | 519 (44) | 497 (83) | 513 (54) |
| Sleep onset latency (min) | 26 (17) | 28 (25) | 20 (23) | 27 (43) | 38 (44) | 18 (11) | 22 (39) | 36 (47) | 24 (29) | 22 (20) | 31 (27) | 17 (9) |
| Sleep efficiency (%) | 86 (7) | 83 (9) | 85 (9) | 86 (9) | 84 (11) | 86 (9) | 86 (10) | 81 (13) | 84 (10) | 86 (7) | 83 (13) | 85 (8) |
| Wake after sleep onset (min) | 51 (39) | 62 (41) | 62 (52) | 48 (32) | 46 (35) | 63 (48) | 50 (49) | 62 (56) | 71 (53) | 59 (44) | 57 (61) | 62 (42) |
| N1 (min) | 28 (21) | 29 (21) | 30 (28) | 30 (26) | 26 (21) | 35 (43) | 31 (22) | 26 (21) | 27 (23) | 34 (25) | 31 (30) | 46 (61) |
| N2 (min) | 266 (63) | 259 (71) | 265 (65) | 249 (62) | 255 (59) | 255 (49) | 270 (50) | 237 (61) | 262 (60) | 266 (39) | 248 (63) | 255 (56) |
| N3 (min) | 99 (30) | 106 (44) | 116 (37) | 106 (37) | 114 (43) | 125 (57) | 104 (27) | 105 (38) | 124 (53) | 99 (30) | 122 (61) | 119 (42) |
| REM (min) | 112 (30) | 83 (37) | 90 (34) | 109 (40) | 109 (33) | 94 (47) | 110 (32) | 95 (35) | 94 (51) | 121 (28) | 96 (37) | 93 (51) |
| N1 (% of TST) | 5.7<br>(4.4) | 6<br>(3.9) | 6.1<br>(5.8) | 6.2<br>(5.1) | 5.1<br>(3.9) | 7.3<br>(8.9) | 6<br>(4.3) | 5.7<br>(4.5) | 5.6<br>(5) | 6.6<br>(5) | 6.1<br>(5.2) | 9.2<br>(12.4) |
| N2 (% of TST) | 52.5<br>(7.6) | 54.0<br>(9.3) | 52.6<br>(10.7) | 50.2<br>(7.1) | 50.5<br>(8.6) | 50.5<br>(9.1) | 52.4<br>(6.2) | 51.5<br>(5.9) | 51.6<br>(10.6) | 51.1<br>(6.1) | 50.0<br>(9.4) | 49.9<br>(10.4) |
| N3 (% of TST) | 20.2<br>(6.7) | 22.9<br>(9.6) | 23.2<br>(7.2) | 22.1<br>(7.8) | 23<br>(8.8) | 24.2<br>(9.8) | 20.5<br>(5.7) | 23.1<br>(7) | 24.4<br>(9.5) | 19.1<br>(5.3) | 24.9<br>(11.6) | 13.1<br>(8.2) |
| REM (% of TST) | 21.7<br>(4.6) | 17.1<br>(6.6) | 18.1<br>(7.1) | 21.5<br>(6) | 21.5<br>(5.1) | 18.1<br>(8) | 21<br>(4.8) | 19.8<br>(4.5) | 18.5<br>(7.5) | 23.2<br>(4.6) | 19<br>(5.8) | 17.7<br>(8.7) |

*Note.* Sleep efficiency was calculated as the percentage of the sleep period (time between sleep onset and final awakening) spent asleep. All values displayed are mean (standard deviation). TST = total sleep time, HC = Healthy controls, FHR = Familial high risk, EC = Early course psychosis.

**Table S3.**
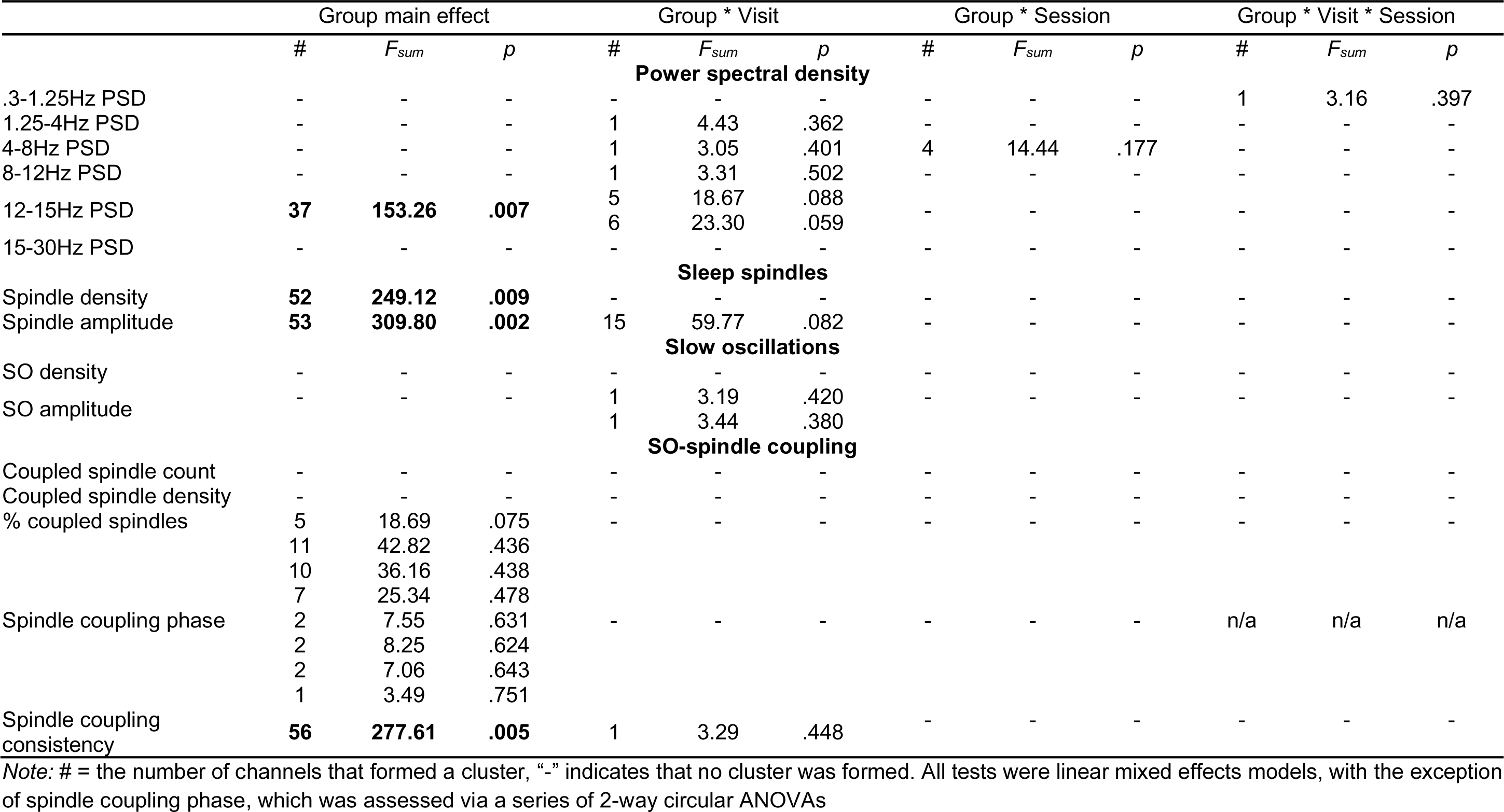
Main effects of group, and interactions involving group, on sleep parameters.

**Figure S1.**
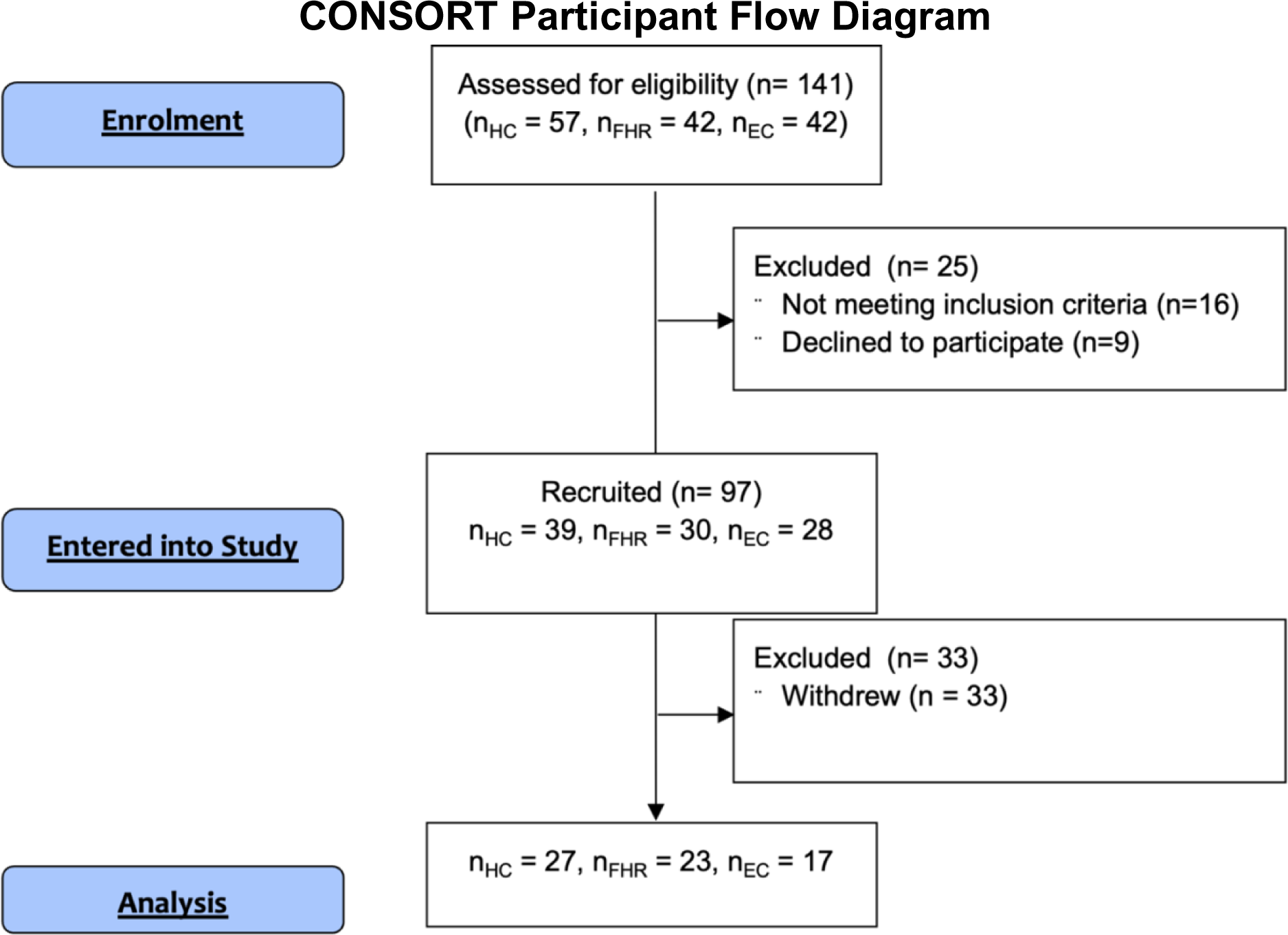
Participant enrolment. HC = Healthy control, FHR = Familial high risk relative, EC = Early course psychosis.

**Figure S2.**
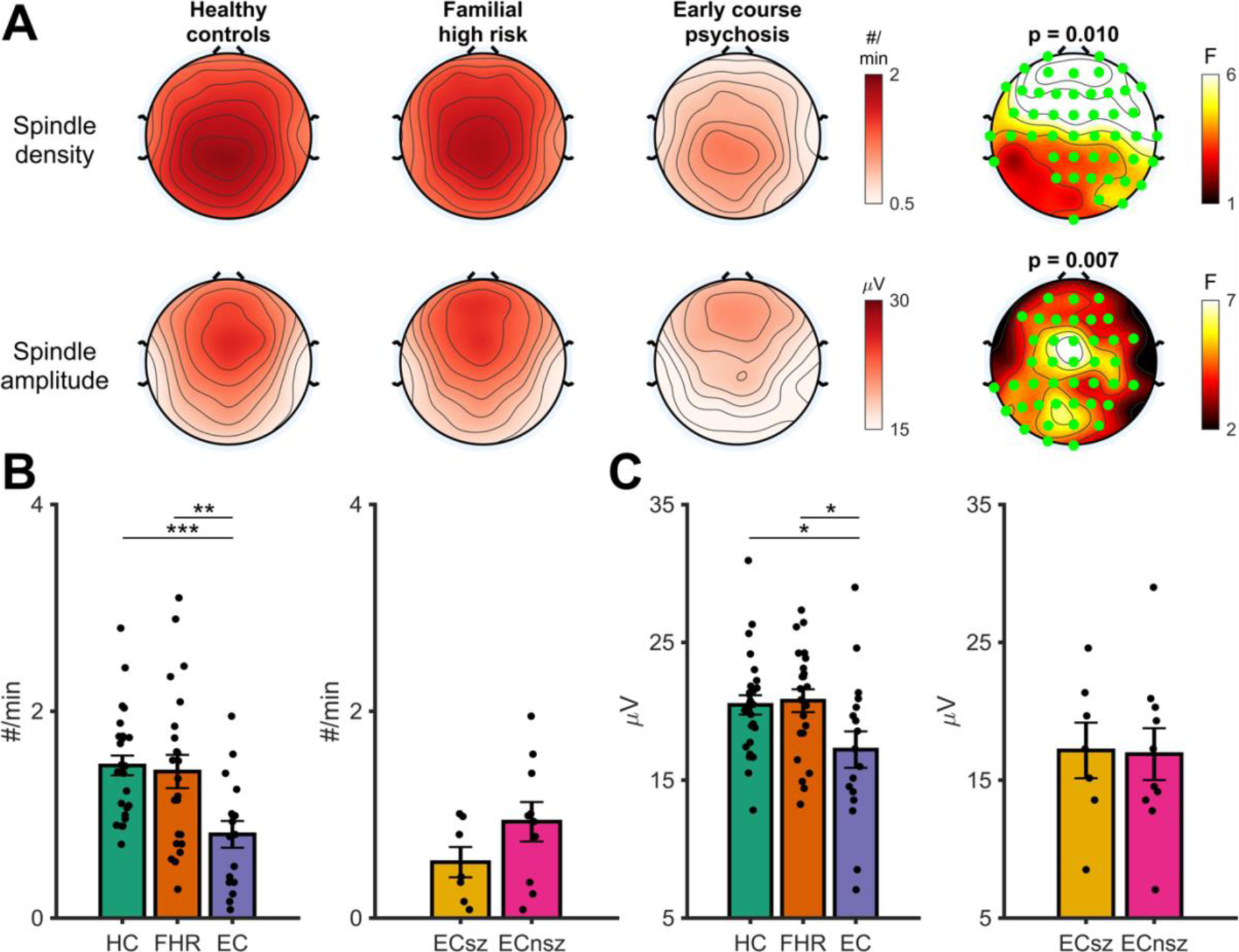
Group differences in N3 sleep spindles. **A** - Topographies showing the main effect of group on N3 spindle density (Row 1) and amplitude (Row 2). N3 spindle topographies (averaged across all four nights) are shown for the three groups separately. The right-hand plot shows F values at each electrode for the main effect of group. Significant electrodes (cluster-corrected) are highlighted in green. Cluster *p-*value displayed above plot. **B** - Pairwise tests for the main effect of group on N3 spindle density with spindle density averaged over significant electrodes in the cluster highlighted in A. Left hand plot shows differences between the three groups. Right hand plot shows differences between schizophrenia and non-schizophrenia psychosis patients. **C** - Same as B, but for N3 spindle amplitude. HC = Healthy controls, FHR = First degree relatives, EC = Early course psychosis, ECsz = Early course schizophrenia, ECnsz = Early course non-schizophrenia psychosis. Error bars indicate the standard error. *** = *p* < .001, ** = *p* < .01, * = *p* < .05 from pairwise estimated marginal means tests.

**Figure S3.**
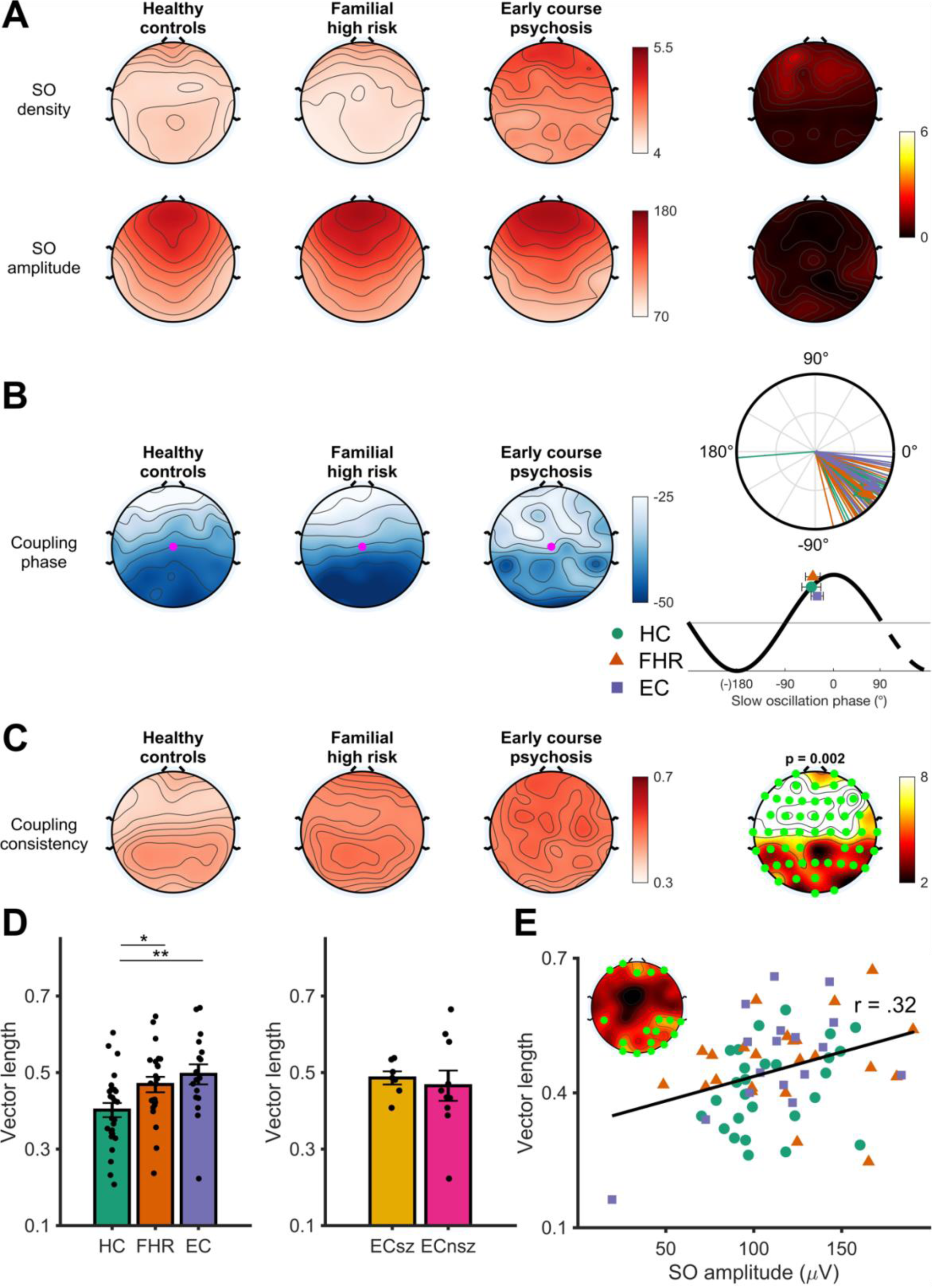
N3 slow oscillation spindle coupling. **A** - Slow oscillation density (top row) and peak-to-peak amplitude (bottom row) in each group, averaged across the four nights. Right most topoplots display F values for the main effect of group. **B** - Preferred coupling phase of spindles to slow oscillations (SOs) in each group, averaged across the four nights. Right: circular phase plot displaying phase distributions across participants for each group at electrode Cz (highlighted electrode in topography). Each line indicates the preferred coupling phase of individual participants. The direction of the arrow indicates the average phase across participants, separately for each group, with the length of the arrow indicating the coupling consistency across participants. A coupling of phase of 0° indicates preferential coupling of spindles at the positive peak of the slow oscillation. A coupling phase of 180° indicates preferential spindle coupling at the negative trough of the slow oscillation. Mapping of SO phase to topographical and circular plots illustrated underneath circular phase plot **C** - Topographies showing the main effect of group on N3 coupling consistency. Coupling consistency (measured as the mean vector length) topographies (averaged across all four nights) are shown for the three groups separately. The right hand plot shows F values at each electrode for the main effect of group. Significant electrodes (cluster-corrected) are highlighted in pink. Cluster *p* value displayed above plot. **D** - Pairwise tests for the main effect of group on N3 coupling consistency, with coupling consistency averaged over significant electrodes in cluster. Left hand plot shows group differences between the three groups. Right hand plot shows differences between schizophrenia and non-schizophrenia psychosis patients. HC = Healthy controls, FHR = First degree relatives, EC = Early course psychosis, ECsz = Early course schizophrenia, ECnsz = Early course non-schizophrenia psychosis. Error bars indicate the standard error. ** = *p* < .01, * = *p* < .05 from pairwise estimated marginal means tests. **E** – Robust linear regression between slow oscillation amplitude and spindle coupling consistency. Insert shows significant electrodes.

**Figure S4.**
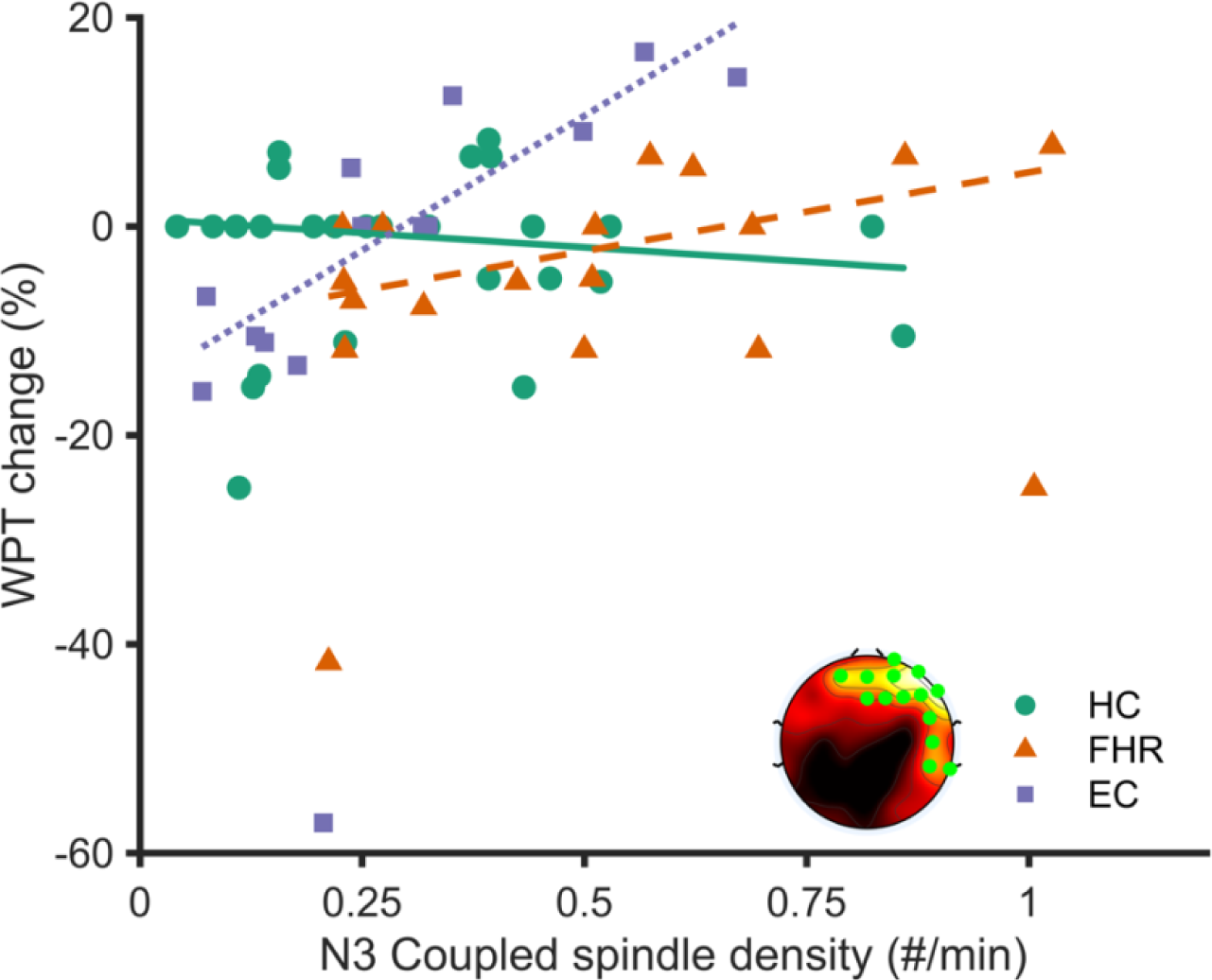
Robust linear regression showing the relationship between coupled N3 sleep spindles and overnight change in word pair memory. Insert shows significant electrodes in the spindle * group interaction (cluster corrected).

